# Subtyping of Small Cell Lung Cancer using plasma cell-free nucleosomes

**DOI:** 10.1101/2022.06.24.497386

**Authors:** Gavriel Fialkoff, Nobuyuki Takahashi, Israa Sharkia, Jenia Gutin, Nadav Hermoni, Rajesh Kumar, Lorinc Pongor, Samantha Nichols, Linda Sciuto, Chante Graham, Parth Desai, Micheal Nirula, Priya Suresh, Melissa Abel, Rajaa Elmeskini, Miriam Maoz, Yakir Rottenberg, Nevo Shoshan, Hovav Nechushtan, Tamar Peretz, Diana Roame, Paula Carter, Ayala Hubert, Jonathan E Cohen, Azzam Salah, Mark Temper, Albert Grinshpun, Zoe Weaver-Ohler, Arun Rajan, William Douglas Figg, Aviad Zick, Ronen Sadeh, Nir Friedman, Anish Thomas

## Abstract

Emerging data on small cell lung cancer (SCLC), an aggressive malignancy with exceptionally poor prognosis, support subtypes driven by distinct transcription regulators, which engender unique therapeutic vulnerabilities. However, the translational potential of these observations is limited by access to tumor biopsies. Here, we leverage chromatin immunoprecipitation of cell-free nucleosomes carrying active chromatin modifications followed by sequencing (cfChIP-seq) on 442 plasma samples from individuals with advanced SCLC, neuroendocrine carcinomas (NEC), non-SCLC cancers, and healthy adults. Beyond providing reliable estimates of SCLC circulating free DNA tumor fraction, cfChIP-seq captures the unique epigenetic states of SCLC tissue- and cell-of-origin. Comparison of cfChIP-seq signals to matched tumor transcriptomes reveals genome-wide concordance, establishing a direct link between gene expression in the tumor and plasma cell-free nucleosomes. Exploiting this link, we develop a classifier that discriminates between SCLC lineage-defining transcription factor subtypes based on cfChIP-seq data. This work sets the stage to non-invasively profile SCLC transcriptomes using plasma cfDNA histone modifications.

## Introduction

Small cell lung cancer (SCLC) is a neuroendocrine lung cancer that is highly aggressive with dismal prognosis, accounting for approximately 15% of all lung cancers^1^. SCLC is one of the solid tumors that sheds the largest amount of cfDNA^2,3^. Prior studies have identified cfDNA mutations in more than 80% of SCLC patients^4–13^, but recurrent targetable mutations in known oncogenes, such as those seen in the kinases that comprise targetable drivers in lung adenocarcinoma, are rare in SCLC^14,15^. Recurrent mutations also do not demonstrate consistent co-occurrence or mutual exclusivity, and thus do not define SCLC subtypes.

SCLCs exhibit high expression of neuronal and neuroendocrine transcription factors and MYC paralogs that drive expression of a broad range of genes related to cell proliferation and growth signaling^14,16^. Importantly, SCLC subtypes driven by distinct transcription factors have unique therapeutic vulnerabilities^17–22^. However, identification of SCLC transcriptomic subtypes and their application in the context of subtype-specific therapies has proven challenging due to limited access to tumor specimens. The majority of SCLC patients do not undergo surgical resection as their disease is detected after it has spread beyond the primary site^23^. Moreover, patients with relapsed disease generally deteriorate quickly, and recurrence suspected on imaging is typically followed by immediate treatment without biopsies. Highlighting this challenge, SCLC is represented in none of the large sequencing initiatives like The Cancer Genome Atlas and Pan-cancer Analysis of Whole Genomes^24^.

Identifying tumor-specific alterations in cell free DNA (cfDNA) presents a powerful opportunity to reduce cancer morbidity and mortality^25,26^. Most of the current clinical applications of cfDNA are centered around interrogating the mutational landscape, and as such are of limited utility in defining transcriptomic subtypes. We recently reported chromatin immunoprecipitation and sequencing of cell-free nucleosomes from human plasma (cfChIP-seq) to infer the transcriptional programs by genome-wide mapping of plasma cell free-nucleosomes carrying specific histone modifications^27^. Specifically, tri-methylation of histone 3 lysine 4 (H3K4me3) is a well characterized histone modification, marking transcription start sites (TSS) of genes that are poised or actively transcribed, and predictive of gene expression^27–30^.

The translational potential of the newly described SCLC transcriptional phenotypes and their associated vulnerabilities is limited by access to tumor biopsies. We hypothesized that plasma histone modifications may non-invasively track tumor gene expression programs, including lineage-defining transcription factors, opening opportunities for subtype-directed therapies for SCLC, currently treated as a single disease. To this end, we applied cfChIP-seq on 305 plasma samples from self-reported healthy subjects (n=33), patients with non-small cell lung cancer (NSCLC, n=16), colorectal cancer (CRC, n=40), neuroendocrine carcinomas (NEC) (n=17) and SCLC (n=67) who had 119 plasma samples collected at multiple timepoints during their treatment. The plasma-based gene expression programs were benchmarked against corresponding tumor transcriptomes, chromatin accessibility, and gene expression assessed by RNA-seq, ATAC-seq, and immunohistochemistry respectively (**Fig. 1A; Supplementary Table 1-2**). Our findings reveal the significant correspondence of plasma cfChIP-seq profiles with tumor gene expression and chromatin accessibility and identify key variables that impact this relationship. By focusing on genes that define SCLC subtypes, we show that plasma cfChIP-seq can effectively classify these subtypes. To confirm these findings, we expanded our analysis to an additional SCLC and NEC validation cohort (n=76/61 plasma samples), further corroborating the potential of plasma histone modifications as biomarkers for tracking tumor gene expression and informing subtype-specific therapies in SCLC.

**Fig. 1:**
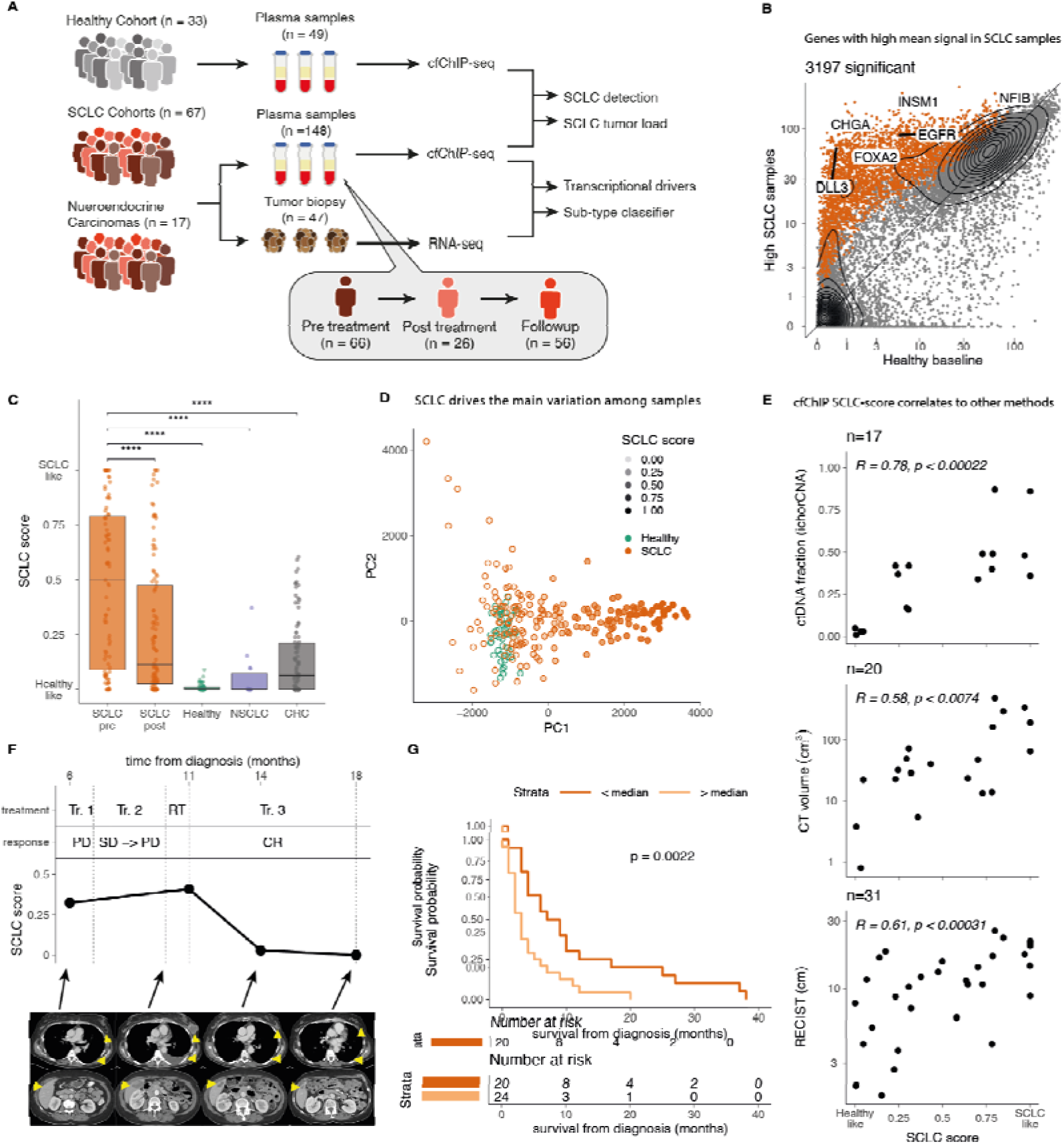
SCLC plasma samples exhibit distinct cfChIP-seq signals that correlate with tumor burden and survival. A. Study outline. Plasma and tumor samples were collected from a cohort of healthy individuals and patients with SCLC at various time points during treatment. cfChIP-seq was performed on plasma samples and RNA-seq on tumor biopsy samples. Plasma cfChIP-seq-based gene expression profiles were benchmarked against tumor RNA-seq, and key SCLC transcription regulators were examined. B. Genes with significantly high coverage in selected SCLC (n=20) samples (Methods) compared to healthy baseline. For each gene, we compare mean normalized coverage in the SCLC samples (y-axis) to a healthy cohort reference (x-axis). 3197 genes with significantly higher coverage (q<0.001) in at least 6 SCLC samples are marked (**Supplementary Table 4)**. C. Median SCLC-score in different cohorts calculated by linear regression (Methods). Each dot is a sample, and boxplots summarize each group’s SCLC-scores distribution. Pre/post refers to the timing of the collection relative to treatment. ****: P < 0.0001. D. Principal components 1 and 2 of SCLC (orange) and healthy (green) samples. Transparency indicates SCLC-score as in C. Principal component analysis was done using all Refseq genes (∼25,000). E. Correlation of SCLC-score with other plasma and imaging-based measures of tumor burden. Abbreviations: RECIST: Response Evaluation Criteria in Solid Tumors. F. Changes of SCLC-score and radiological tumor burden in a patient with SCLC over the treatment time-course (SCLC0191). Abbreviations: Tr. 1 - durvalumab & olaparib; Tr. 2 - Topotecan & M6620; RT - radiation; Tr. 3 - investigational therapy; CR - complete response; SD - stable disease; PD - progressive disease. G. Kaplan-Meier survival plot in patients with SCLC-score above (red; median survival: 3 month) or below (blue; median survival: 8.5 month) median. The survival curve was conducted on pre-treatment samples. *p* calculated by log-rank test.

## Results

### SCLCs have distinct cfChIP-seq signals that track tumor burden and prognosis

Plasma samples were collected, processed, and H3K4me3 ChIP-seq^27^ performed directly on ∼1ml of plasma (Methods) with a median of 3.5 million unique reads sequenced per sample (**Supplementary Table 3**). The number of normalized reads mapping to its respective TSS region(s) was computed for every gene, resulting in gene counts analogous to transcription counts in RNA-seq data. Comparing the gene counts in SCLC plasma samples to those in healthy reference samples (Methods), we found significantly elevated counts in hundreds to thousands of genes (**Fig. 1B, S1A**).

cfDNA of cancer patients consists of DNA fragments originating from tumor cells and DNA released by normal cells, primarily from the hematopoietic lineage^27,31^. The tumor-derived fraction of cfDNA can vary substantially, depending on several variables including tumor burden and growth activity^32^. To account for this variability, we developed an ‘SCLC-score’ that reflects the proportion of tumor-derived cfDNA. The score is generated using a linear regression model that matches the observed gene count profile in a sample to a weighted mixture of reference cfChIP-seq profiles. The reference dataset consisted of plasma samples from healthy individuals and a representative SCLC archetype derived from the 10% of SCLC samples (n = 20), with the highest number of differential genes compared to healthy samples (**Fig. S1A**, Methods).

The SCLC-score ranged from 0 (indicating a ‘healthy like’ profile) to 1 (‘SCLC like’). In SCLC patient samples collected before treatment, the median SCLC-score was 0.5, which dropped to 0.11 post-treatment, suggesting a reduction in tumor-derived cfDNA due to therapy. In contrast, plasma from healthy subjects and patients with non-SCLC cancers displayed absent or very low SCLC-scores (median of 0 in healthy and NSCLC, 0.06 in CRC; ANOVA p < 10^-15^. **Fig. 1C**). To validate our estimation in an unsupervised manner, we performed principal component analysis (PCA) on the gene counts of healthy controls and SCLC samples (Methods). When we mapped the SCLC-score onto the two-dimensional PCA plot, PCA1 - the axis explaining the most sample variability - showed a near-perfect correlation with the estimated tumor load (*r* = 0.98, p < 10^-15^ **Fig. 1D, S1B-C**).

Importantly, cfChIP-seq SCLC-scores were significantly correlated with multiple other measures of tumor fraction, including somatic copy number alteration-based estimates from ultra-low pass whole genome sequencing^33^, circulating tumor cell (CTC) counts, total cfDNA concentrations (Methods. Pearson r = 0.78, 0.43 and 0.61; p < 2×10^-4^, 0.01, and 0.03, respectively), computerized tomography scan-based volumetric tumor assessments, and standardized unidimensional tumor measurements^34^ (Pearson r and p: 0.58 and 0.61; <0.007 and <3×10^-4^, respectively **Fig. 1E** and **S1D-E**). Furthermore, cfChIP-seq SCLC-scores tracked radiographic tumor burden through the treatment time course (**Fig. 1F**), and predicted treatment response and overall survival (**Fig. 1G** and **S1F-G**).

Taken together, these results demonstrate the potential of cfChIP-seq to non-invasively detect and quantify SCLC from plasma. Further investigations are warranted to explore whether this approach could be adapted for lung cancer screening, with the prospect of significantly improving patient survival through early detection.

### cfChIP-seq recovers SCLC tissue and cellular origins

We next investigated whether plasma cfChIP-seq signals could provide insights into the epigenetic state of tissue and cells of origin of SCLC tumors. Among the more than 3,500 genes with significantly elevated coverage in SCLC plasma (**Fig. 1B**, methods) many were specifically elevated in SCLC samples compared with healthy controls or other cancers. Interestingly, the signal across these genes generalized to plasma samples from patients with NECs. These tumors can arise at almost any anatomic site, and exhibit tissue-independent convergence to a neuroendocrine histology, while maintaining molecular divergence driven by distinct transcriptional regulators<u>(Wang et al. 2024)</u>. These results underline the similarities in the transcription patterns of the SCLC and NEC. (**Fig. 2A-B**). Notably, these SCLC-signature genes showed a remarkable enrichment for genes expressed in SCLC cell lines^35^ and pulmonary neuroendocrine cells^36^, with significant overlaps of 272 out of 465 and 53 out of 92 genes (p <7.8*10^-97^ and <3.5*10^-15^, respectively)^37,38^. Specifically, SCLC plasma exhibited elevated counts of canonical SCLC genes such as DLL3, INSM1, CHGA, and CRMP1, with significantly higher levels than those observed in healthy samples (**Fig. 2C-D**).

**Fig. 2:**
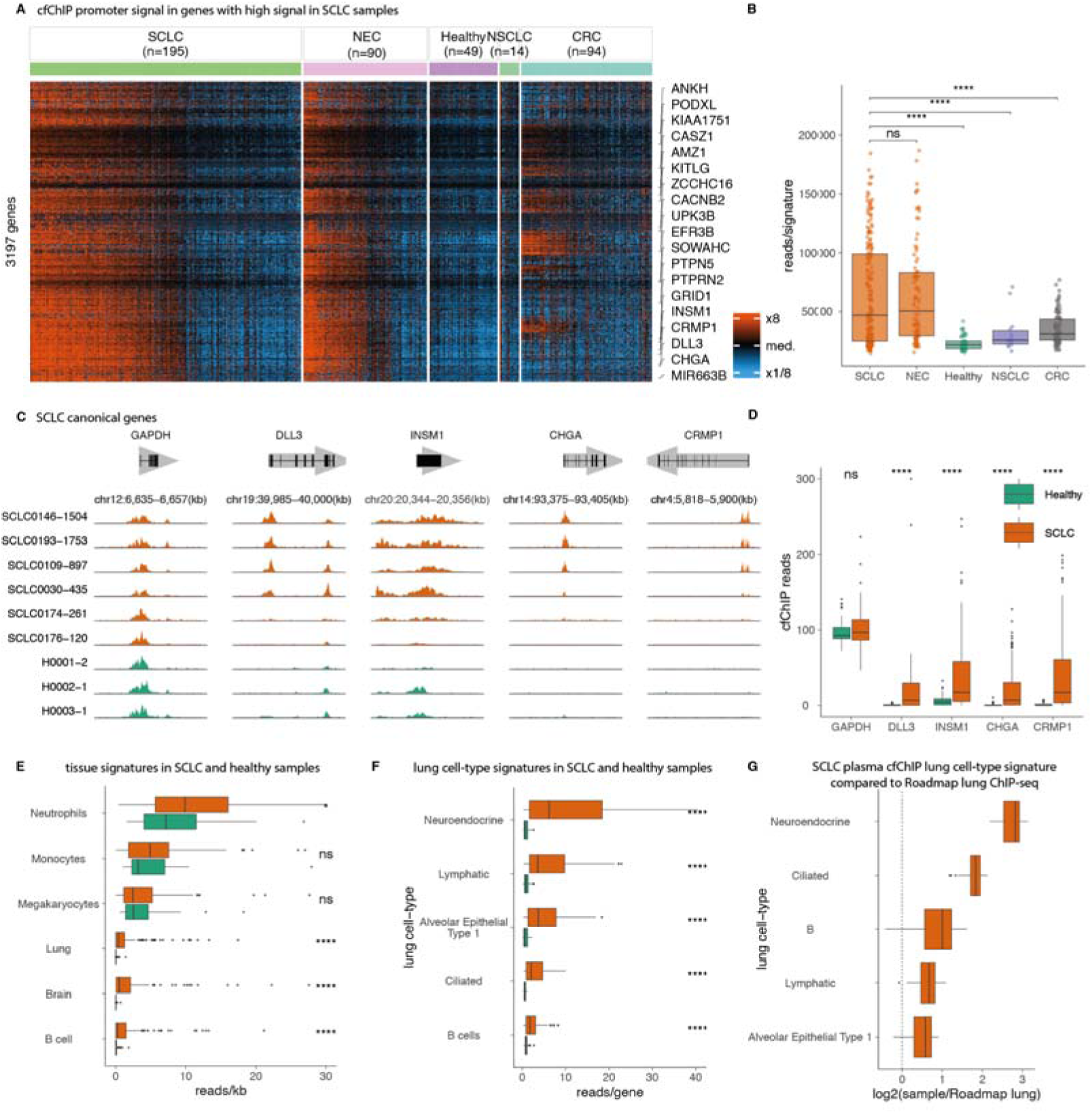
cfChIP-seq recovers SCLC tissue and cellular origins. A. Hierarchical clustering of SCLC signature genes (rows) across the study cohorts (columns). Samples within each cohort are ordered by the SCLC score. Normalized coverage on the gene promoter was log-transformed (log2(1+coverage)) and adjusted to zero mean for each gene across the samples. Abbreviations: SCLC, small cell lung cancer; NEC, neuroendocrine cancer; NSCLC, non-small cell lung cancer; CRC, colorectal cancer. B. Median and distribution of coverage on genes shown in A across the study cohorts. ****: P < 0.0001 C. Genome browser view of cfChIP-seq signal in canonical SCLC genes (*DLL3*, *INSM1, CHGA, CRMP1*) and *GAPDH* as control. Orange and green tracks represent SCLC and healthy samples respectively. D. Median and distribution of the cumulative cfChIP-seq coverage over the SCLC-signature genes. E. Cell and tissue of origin signatures in healthy and SCLC samples. x-axis indicates the absolute contribution of signature (normalized reads/kb corrected by estimated cfDNA concentration; methods) F. Single-cell RNA-seq-derived lung cell-type signatures in healthy and SCLC samples. x-axis indicates the sum of normalized reads in the marker genes of every tissue. Ratio for signal in cell-types shown in F of high-score SCLC samples (n=23) compared to Encode3 lung ChIP-seq.

To characterize the tissue-specific origins of cfDNA, we established a set of tissue-specific genomic loci using ChIP-seq reference data^39,40^ (Methods). Using this approach, we confirmed that the signals in healthy plasma are mainly derived from neutrophils, megakaryocytes, and monocytes^27,31^. In contrast, signals in SCLC plasma exhibited additional signals derived from cells with characteristics of lung and brain tissues, as well as B-cells, suggesting contributions from cells of pulmonary, neuronal, and lymphocyte lineages (**Fig. 2E** and **S2A**). These tissue-specific signals positively correlated with the SCLC-score, reinforcing their relevance (**Fig. S2B**).

To further investigate whether cfChIP-seq can provide clues to the cell-of-origin, we applied lung cell-type-specific signatures derived from single-cell RNA-seq data ^41^. Across the ∼50 cell-types examined, spanning lung epithelial, endothelial, stromal, and immune cells, we observed a strong enrichment of neuroendocrine cell type in SCLC plasma compared to healthy controls (**Fig. 2F**). This finding is especially striking since neuroendocrine cells constitute only 0.13% of a healthy lung tissue^41^. Furthermore, the neuroendocrine cell signature was significantly elevated in SCLC plasma relative to normal lung tissue (**Fig. 2G**). Signals of other lung cells including ciliated cells and alveolar epithelial type 1 cells were also higher in SCLC plasma compared to healthy controls (**Fig. 2F**, **S2C** and **S2D**), hinting at the possibility of SCLCs arising additionally from non-neuroendocrine cells-of-origin as previously described^42^ or indicating injury to the specific cells.

Together, these findings demonstrate that cfChIP-seq can capture the unique epigenetic states of tissues and cells-of-origin associated with SCLC tumors.

### Plasma cfChIP-seq reflects tumor chromatin accessibility state

Since chromatin accessibility is closely linked to gene expression programs, we asked whether circulating nucleosomes reflect the chromatin accessibility state of the corresponding tumors. Due to the challenges of mapping chromatin accessibility in small biopsy samples, we generated patient-derived xenografts (PDX) (**Fig. 3A**) from tumors of patients who had plasma profiled using cfChIP-seq, and performed transposase-accessible chromatin using sequencing (ATAC-seq)^43^ on these PDX tumors. The relationship between ATAC-seq and the H3K4me3 mark is complementary, as ATAC-seq identifies regions of open chromatin and H3K4me3 marks active promoters.

**Figure 3:**
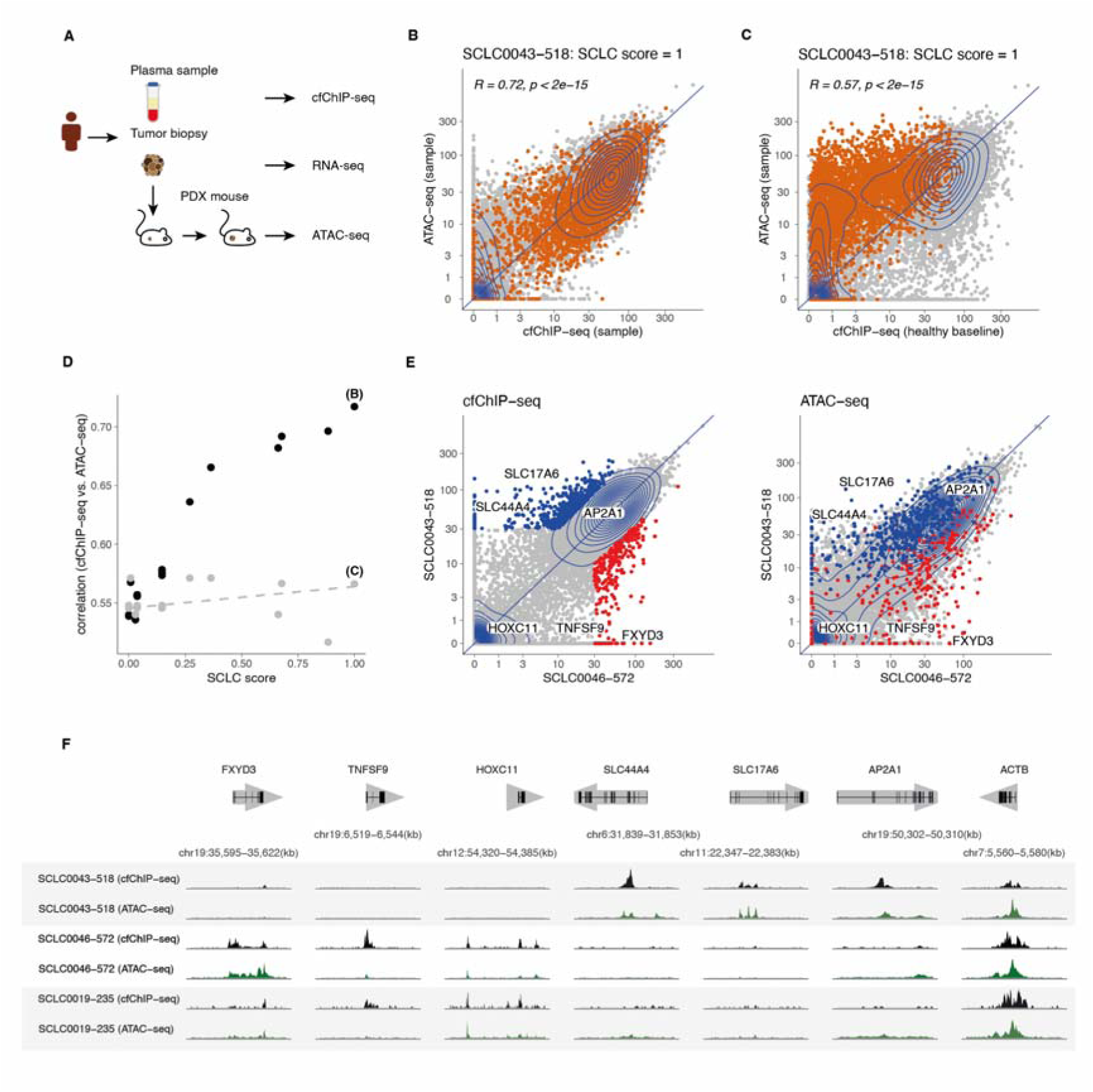
Plasma cfChIP-seq reflects tumor chromatin accessibility state. A. Schematic outline of ATAC-seq experiment. Patient-derived xenograft (PDX) models were established by transplanting tumor tissue obtained directly from patients subcutaneously into immunocompromised mice. The resulting tumor is harvested and subject to ATAC-seq. B. Comparison of ATAC-seq (y-axis) and cfChIP-seq (x-axis) promoter coverage of patient-matched samples over all Refseq genes. Every dot represents a gene promoter. Colored dots represent the 3684 SCLC signature genes as in Figure 1B. C. Y-axis is the same as in B. x-axis represents the average cfChIP-seq coverage in healthy individuals (mean coverage of healthy cfChIP-samples). D. Correlation of ATAC-seq and cfChIP-seq promoter coverage of corresponding samples (y-axis) as a function of the SCLC-score (x-axis). The gray dots represent the correlation of the same ATAC-seq samples to the healthy baseline. The labeled dots correspond to panels B and C. E. Comparison of differential genes pattern across cfChIP-seq (right) and the corresponding ATAC-seq (left) data in samples from two different patients. Genes in ATAC-seq (right) are colored by their differential status in the cfChIP-seq (left). F. Genome browser view of examples of genes that show differential signals between the two samples shown in D and a control gene (*ACTB*). Three pairs of cfChIP-seq and the corresponding ATAC-seq are shown.

When comparing the ATAC-seq (PDX) and cfChIP-seq (plasma) promoter coverage in a sample with high plasma SCLC-score, we observed a strong correlation between the two assays (*R* = 0.72, *p* < 2×10^-15^; **Fig. 3B**). In particular, the SCLC signature genes are predominantly aligned along the diagonal. This high correlation is not seen when comparing ATAC-seq coverage to the average cfChIP-seq coverage in the healthy cohort, suggesting that much of the correlation is driven by the SCLC signature genes that exhibit increased ATAC-seq signal and elevated cfChIP-seq signal (**Fig. 3C**). Comparison of ATAC-seq vs. cfChIP-seq across all sample pairs demonstrated that the correlation between the assays was associated with increasing SCLC scores, well beyond what could be attributed to chance (**Fig. 3D**).

To examine whether differences in plasma cfChIP-seq signal between samples correlate with differences in tumor chromatin accessibility, we compared the cfChIP-seq data from two patients with high SCLC scores. Remarkably, the differentially active genes between the two plasma samples exhibited a similar pattern of differential accessibility in the corresponding tumor ATAC-seq data (**Fig. 3E-F**).

Collectively, these findings suggest that plasma nucleosomal H3K4me3 marks can serve as reliable indicators of the tumor chromatin accessibility state.

### Plasma cfChIP-seq informs tumor gene expression

Recent studies reveal that SCLC tumors are transcriptionally heterogeneous^21,44,45^ (**Fig. 4A**). We sought to understand whether plasma cfChIP-seq reflects gene expression patterns of SCLC tumors.

**Fig. 4:**
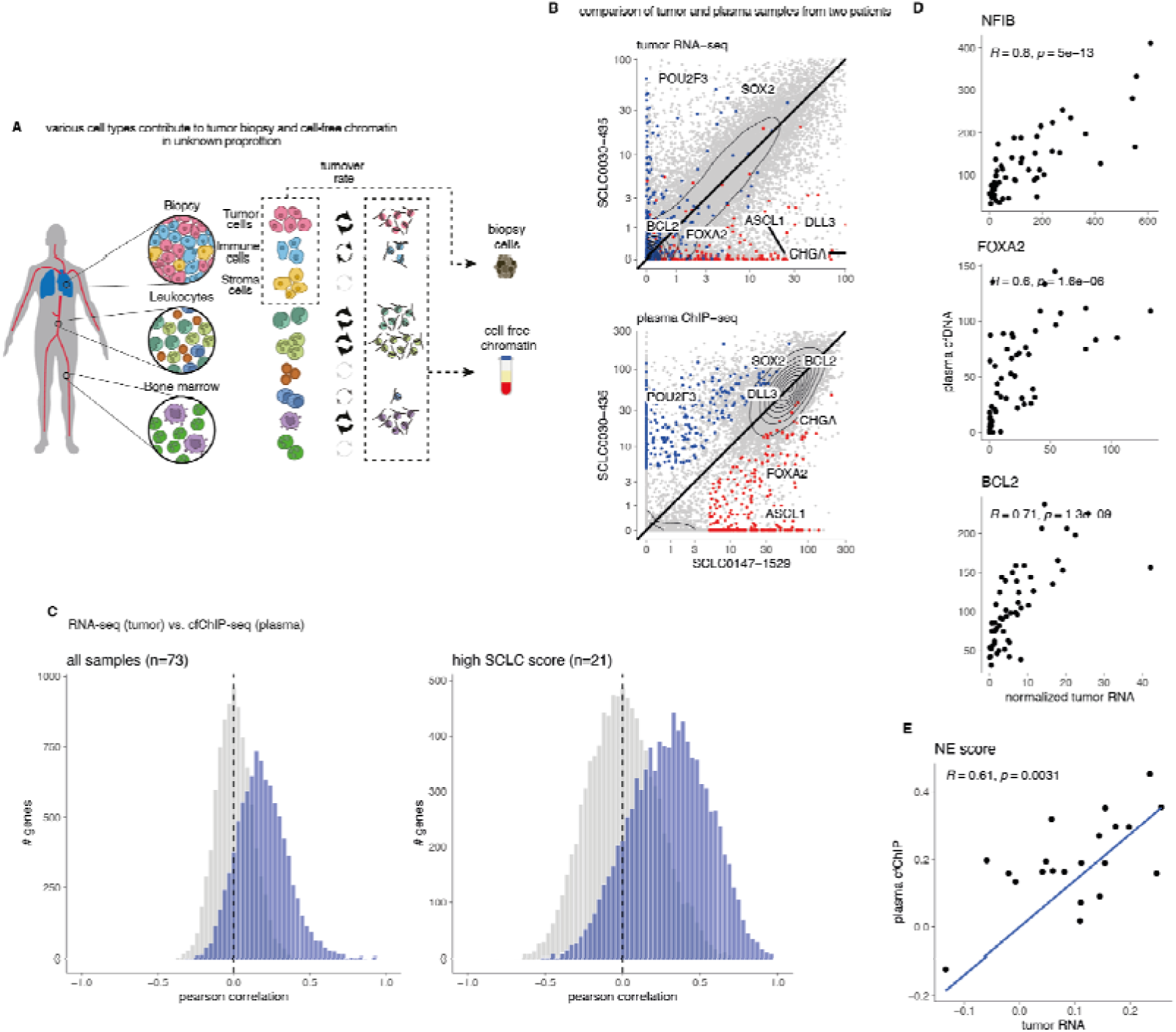
Plasma cfChIP-seq informs tumor gene expression. A. Concept figure illustrating the various sources and proportions of cells sampled in plasma cfChIP-seq and tumor RNA-seq. The data obtained from these two assays is derived from a different unknown mixture of multiple types of cells. B. Comparison of gene expression in two high SCLC-score samples. Top: TMM-normalized CPM of tumor RNA-seq. Bottom: normalized gene counts of cfChIP-seq. blue and red points indicate genes that had high cfChIP-seq gene counts in one sample compared to the other and low gene counts in healthy reference cfChIP. C. Gene level analysis of the correlation between tumor gene expression and plasma cfChIP-seq coverage across individuals with matched tumor and plasma samples (left: all SCLC samples. right: high SCLC-score samples). For each gene, we computed the Pearson correlation of its tumor expression and the normalized cfChIP-seq coverage across the samples. Shown is a histogram of the correlations on genes with high dynamic ranges (Methods). In gray is the histogram of a random permutation of the relation between tumor expression and plasma cfChIP. D. Examples of correlation for several known SCLC oncogenes. E. Correlation of NE-score computed based on plasma cfChIP-seq and tumor RNA-seq. Only plasma samples with matching tumors and high SCLC-score are presented.

As an example, among the high SCLC-score plasma samples, one sample (SCLC030-435) in particular exhibited markedly higher signal at several genes (e.g., *POU2F3* and *BCL2*) compared to another high SCLC-score sample (SCLC147-1529, **Fig. 4B**). Examining time-point matched tumor RNA-seq of these samples revealed differential expression of many of the same genes, indicating that differential cfChIP-seq signals might indeed reflect tumor gene expression (**Fig. 4B** and **S3A**).

To systematically examine the relationship between circulating chromatin state and tumor gene expression, we compared cfChIP-seq profiles and RNA-seq data from the same patients. Ideally, we would want time-matched samples where both report on the same tumor state. However, due to clinical constraints, some comparisons involved RNA-seq from an earlier biopsy and cfChIP-seq at the time of disease recurrence (**Supplementary Table 1**, **Fig. S3B**). To address this, we performed analyses on both time-matched samples (n = 36; smaller numbers but better correspondence) and the full set of samples (n = 73; larger numbers but potential for weaker correspondence). Additionally, since the tumor’s contribution to cfDNA may be low in some cases, we focused on high SCLC-score samples while also considering all available samples.

We computed for each gene the correlation between plasma cfChIP-seq counts and tumor RNA-seq TMM-normalized CPM values. Excluding genes with low dynamic range in cfChIP-seq or RNA-seq (Methods), a significant positive correlation was observed in more than 25% of the genes (2286 genes with q < 0.05; Pearson 0.29 < *r* < 0.97) of the full set of samples. When comparing only the high SCLC-score samples with matched tumor sample (n=21), despite the reduced statistical power, we still observed a large number of significant genes (16%, 1524 genes with *q* < 0.05, 931 of them overlapping with the significant genes in the full dataset) showing higher correlation (Pearson 0.56 < *r* < 0.99. **Fig. 4C** and **S4A**). In particular, a high positive correlation was observed between cfChIP-seq and RNA-seq read counts for several important SCLC oncogenes such as *BCL2*, *NFIB*, and *SOX2* (**Fig. 4D**).

The agreement at the level of genes translates to the agreement of groups of genes. As an example, we used a SCLC neuroendocrine status gene signature^46^. We removed from this set genes that are high in healthy plasma samples (Methods) and compared the cfChIP-seq aggregated signal over these genes to the aggregated RNA-seq of these genes across samples with high tumor load (n = 21). This analysis revealed a significant positive correlation (**Fig. 4E**; Pearson *r =* 0.75 and *p* < 0.0001).

To better understand the concordance between tumor RNA-seq and cfChIP-seq, we examined factors contributing to variations in correlation. One source of discordance can be poised promoters where the chromatin is accessible and marked by H3K4me3, but the gene is transcriptionally inactive. In such instances, the cfChIP-seq signal may exhibit varying signals, whereas the corresponding RNA-seq data would show no signal. Additionally, chromatin marks are binary at the single-cell level (either present or absent), whereas RNA levels span a broader dynamic range. Notebly, genes displaying a cfChIP-seq high dynamic range tend to have stronger correlation (**Fig. S4B**). Another potential source divergence between the cfChIP-seq and tumor RNA-seq stems from the contribution of non-tumorous tissues to each measurement (**Fig. 4A** and **Fig. S3C-E**, **Supplementary Table 6**).

Overall, these findings demonstrate that cfDNA chromatin state, as assessed by cfChIP-seq, informs the tumor gene expression programs, especially in plasma samples with high tumor fraction. Moreover, these findings highlight crucial variables that influence the concordance between cfDNA chromatin state and tumor gene expression.

### Plasma cfChIP-seq predicts tumor gene expression of SCLC lineage-defining transcription factors

Building on the concordance between tumor gene expression and plasma chromatin state, we next focused on key transcription factors central to SCLC tumorigenesis. Transcriptional signatures of SCLC heterogeneity converge on two major cell states, namely neuroendocrine (NE) and non-neuroendocrine (non-NE)^46,47^. The SCLC cell states are further characterized by the expression of key lineage-defining transcription factors, ASCL1 and NEUROD1 defining NE cell states, and POU2F3 defining non-NE cell states. A fourth subgroup has been characterized by expression of YAP1^48,49^ or low expression of all three transcription factors, accompanied by an inflamed gene signature^19^.

Most tumors in our cohort had high expression of NE-lineage defining genes *ASCL1* and *NEUROD1*, with co-expression of both in some cases (**Fig. 5A**). *POU2F3*, *YAP1*, and a newly described subtype marker ATOH1^1,17,50^ were expressed less frequently.

**Fig. 5:**
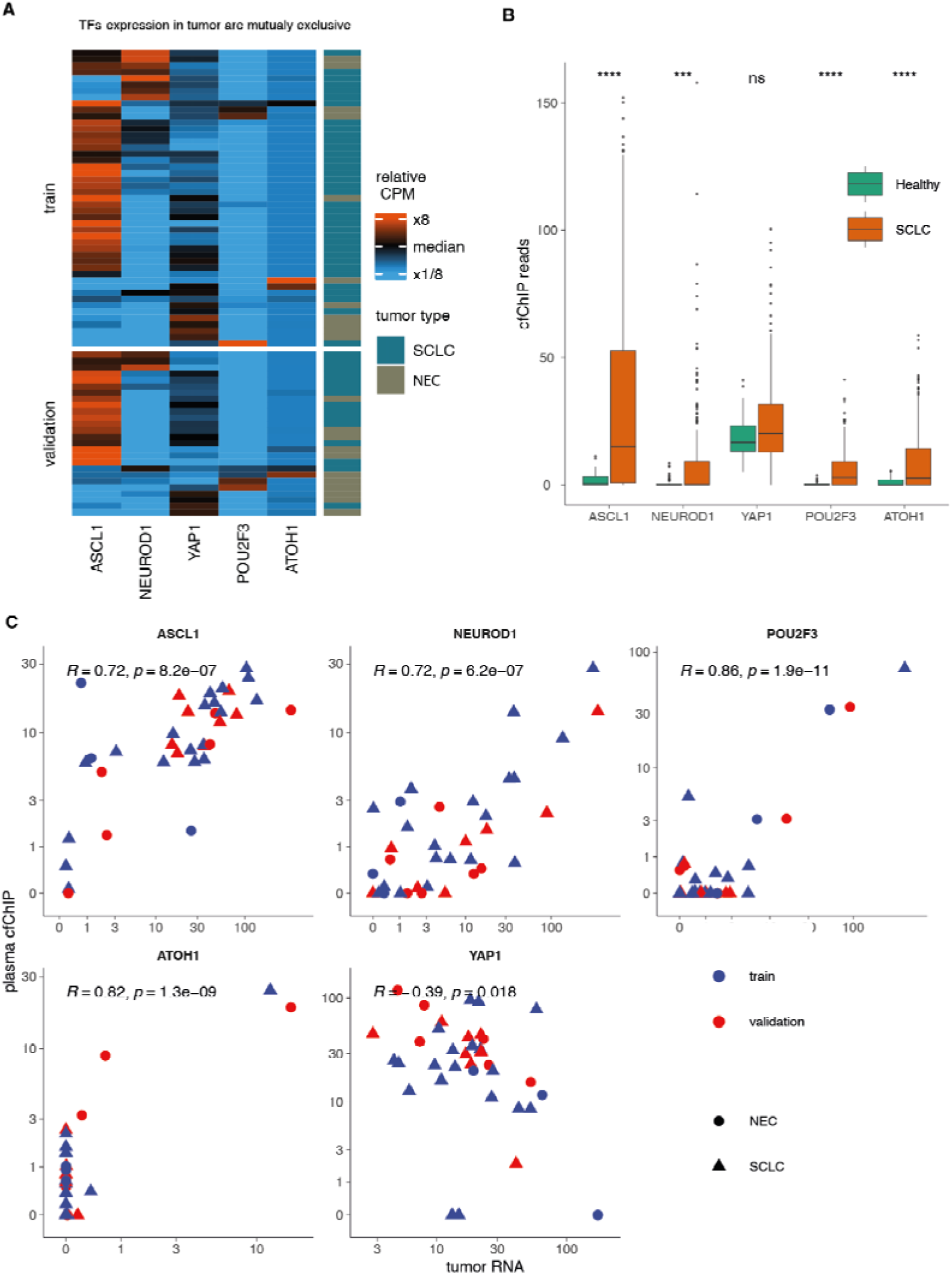
cfChIP-seq displays differential expression of SCLC transcription drivers. A. Heatmap showing relative expression levels of 5 canonical SCLC transcription drivers across the tumor RNA-seq samples. Values are presented in TMM-normalized CPM and adjusted to zero mean for each sample across the 5 genes. Transcription drivers’ expression patterns are generally mutually exclusive. B. Median and distribution of SCLC and healthy cfChIP-seq plasma sample coverage on the gene shown in A. **** *p* < 0.0001. C. Correlation of plasma cfChIP-seq coverage and tumor RNA-seq of the genes shown in C. Only plasma samples with high SCLC-score (n=36) are presented. Color indicated the train/validation status of the sample and shape represents the tumor type.

We sought to examine whether the expression of SCLC lineage-defining transcription factors in the tumor is reflected in the plasma cfChIP-seq. cfChIP-seq counts for *ASCL1*, *NEUROD1*, and *POU2F3* were significantly elevated in SCLC plasma, compared to barely detectable levels in healthy plasma. However, *YAP1* counts were similar among healthy and SCLC plasma samples (**Fig. 5B** and **S5A**), likely due to *YAP1* activity in normal tissue contributing to cfDNA, as suggested by H3K4me3 marks on *YAP1* promoters in normal tissue^39^.

To evaluate whether these cfChIP-seq signals accurately reflect tumor gene expression in individual patients, we assessed their correlation with RNA-seq data from tumor samples in high SCLC-score cases (n=36). A strong correlation between cfChIP-seq and tumor RNA-seq was observed for *ASCL1*, *NEUROD1*, and *POU2F3* (Pearson *r* = 0.72, 0.72, and 0.86; p < 8.2×10^-7^,6.2×10^-7^, and 1.9×10^-11^ respectively), all of which which were absent from healthy control cfDNA. A similarly high positive correlation was observed for *ATOH1* (Pearson r = 0.82; p < 1.3×10^-9^). This correlation was observed both in the SCLC and NEC samples (**Fig. 5C**). Notably, while the TSS of *POU2F3* and *ATOH1* were marked by H3K4me3 in many of the SCLC samples, only in a subset of them did the signals span beyond the TSS region to a wider region of approximately 10KB, suggesting that in these cases they are involved in cell-type-specific functions^51^. We find these additional regions correlated best with gene expression in the tumor (**Fig. S5B-C**).

To validate the agreement between cfChIP-seq signals and tumor gene expression at the protein level, we performed immunohistochemistry of INSM1 on SCLC tumors (n=10).

INSM1 is a standard marker used in diagnostic pathology of SCLC, and a super-enhancer-associated transcription factor that regulates global neuroendocrine gene expression ^52^. The results revealed high concordance between plasma cfChIP-seq and tumor protein levels (*r* = 0.74, *p* = 0.014; **Fig. S5D**).

Overall, these results demonstrate that plasma cfChIP-seq can reliably capture the expression patterns of SCLC lineage-defining transcription factors, offering insight into tumor cell states and heterogeneity.

### Subtyping of SCLC from plasma cfChIP-seq

Next, we sought to explore the possibility of subtyping SCLC tumors directly from plasma using cfChIP-seq. Although SCLC subtyping is often viewed as a discrete classification between the types, we observed cases where the tumors exhibited high levels of two or more lineage-defining transcription factors simultaneously (**Fig. 5B**). This aligns with data from recent studies^55^. Therefore, we decided to predict the activity of each factor separately, examining four distinct classifications for each sample.

A straightforward strategy for classifying the lineage-defining transcription factors subtypes is to classify based on the cfChIP-seq signal strength for each transcription factor. Evaluating the predictive performance of this method using ROC curves, we observed high area-under-the-curve (AUC) scores for *NEUROD1*, *POU2F3* and *ATOH1*, even in samples with SCLC scores as low as 0.05 (**Fig. 6A,B**).

**Figure 6:**
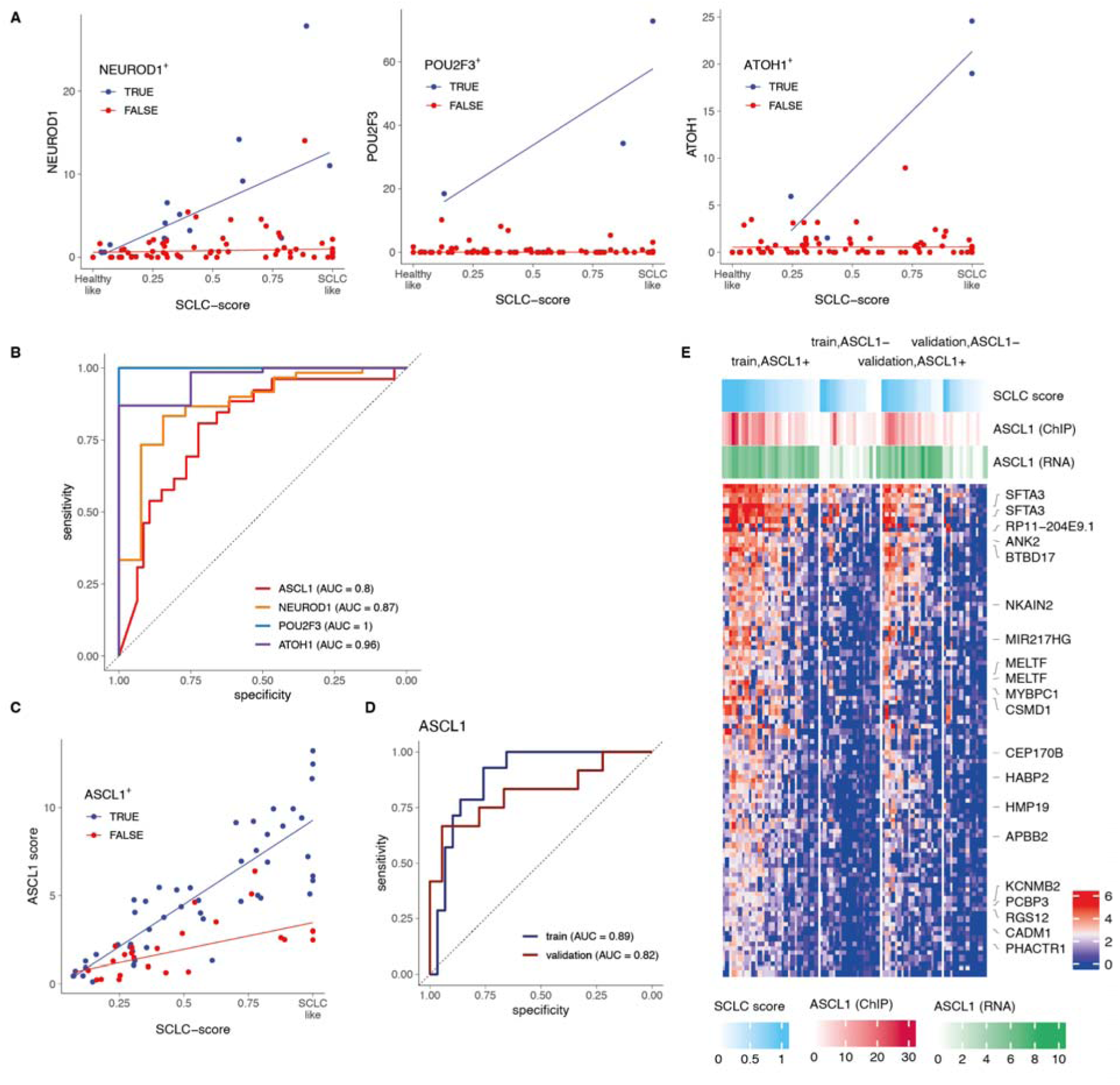
SCLC subtyping using cfChIP-seq signals. A. SCLC score (x-axis) and cfChIP-seq promoter coverage (y-axis) of samples that are annotated as positive (blue) and negative (red) to *NEUROD1*, *POU2F3* and *ATOH1* based on biopsy RNA-seq. Linear fit shows a positive relation between the promoter coverage and SCLC-score across the positive samples while the negative samples remain low and constant. B. ROC plot summarizing the performance of the single-gene classifiers. C. *ASCL1*-specific signature strength (y-axis) increases linearly with SCLC-score (x-axis) in *ASCL1*^high^ samples D. Performance of the *ASCL1* signature score on the training and validation set. Training set values were obtained using leave-patient-out cross-validation. E. cfChIP-seq coverage patterns of *ASCL1-*specific genomic regions (rows) across the training and validation samples (columns). Color scale represents the normalized coverage on the genomic region. Top bars display the ASCL1 cfChIP-seq promoter coverage, ASCL1 score, log2(1+RNA-seq CPM) and SCLC-score values across the samples. The order of samples within each group is determined by the SCLC-score.

However, single-gene-based classification proved challenging for *ASCL1*. A subset of samples displayed a high cfChIP-seq signal at the *ASCL1* promoter but low RNA expression, (**Fig. 5D**), suggesting either a poised but inactive promoter or tumor heterogeneity where the biopsy was not representative of all the tumor. We hypothesized that employing a multigene signature, taking into account genes that are regulated by *ASCL1*, would offer a more robust classifier.

To define such a set of genes, we applied a t-test to genomic regions influenced by the estimated tumor contribution (Methods). We then performed leave-one-patient-out cross-validation on the training data to determine the optimal number of regions for the signature, selecting 100 regions based on performance metrics (**Supplementary Table 7)**. The aggregated read counts over the signature displayed a linear relationship with the SCLC-score in positive samples but remained constant and low in other samples, even those with high SCLC-scores. The ratio of signature counts to SCLC-score effectively discriminated between the positive and negative samples, achieving a high predictive performance (AUC 0.82 on the validation cohort). The genes of the signature were significantly enriched for “neuroendocrine cells in the lung” (p < 10^-4^; Descartes Cell Types and Tissue 2021^37,38^) (**Fig. 6C-E**).

These findings suggest that cfChIP-seq can address an unmet need of molecularly classifying SCLC into transcriptomic subsets in a minimally invasive manner, directly from plasma. This approach could provide valuable insights for clinical decision-making and precision medicine strategies in SCLC.

## Discussion

Cancer master transcription factors control global gene expression programs^17,56–58^, and are attractive targets to classify and treat cancer given their essential role in driving distinctive cell identities^59,60^ and the tendency for cancer cells to become highly dependent on their sustained and high-level expression^61^. However, transcription factor profiling in clinical tumor samples is challenging, particularly for cancers where access to tumor specimens is limited. Recent studies have identified subtypes of SCLC, the most lethal type of lung cancer, defined by expression of lineage-defining transcription factors, and their unique therapeutic vulnerabilities^17–21^. Yet, further testing and broad clinical application of these findings is limited by the availability of tumor tissue, a consequence of the widely metastatic and aggressive nature of SCLC, which typically precludes surgical resection at diagnosis and tumor biopsies at relapse. As such, under current guidelines, all patients with SCLC receive the same treatments. Here, we apply plasma cfChIP-seq which reports the promoter state of cell-free chromatin in circulation^27^ to SCLC samples. We find that cfChIP-seq recovers the unique epigenetic states of SCLC tissue and cell of origin, and importantly tumor gene expression, particularly SCLC lineage-defining transcription factors, providing a systematic view of tumor state, opening the possibility of molecularly classifying SCLC directly from as little as 1 ml of plasma.

Our findings reveal that SCLC has a distinct cell-free chromatin signature, which can be detected in patient plasma using cfChIP-seq. This signature can differentiate SCLC from other cancers and healthy controls and can be detected even when it has low representation in the plasma (e.g., after therapy), and is highly correlated with serologic and radiological estimations of tumor burden and prognosis.

In matched plasma and tumor biopsy samples, we show for the first time, the concordance of gene expression inferred from plasma cell-free chromatin and tumor transcriptome at the level of the individual patient. This concordance opens a new avenue to study the heterogeneity between SCLC tumors, tumor response to treatment, and the transcriptional changes in the tumor throughout the patient trajectory.

Importantly, cfChIP-seq profiles identify the activity of key SCLC transcriptional drivers, including ASCL1 and NEUROD1 that drive NE phenotypes and POU2F3 that drive a non-NE phenotype^19,44,45,47^. These results set the stage for non-invasive subtyping and molecular profile-based treatments for patients with SCLC, which might be more effective than the current one-size fits all approach. A larger patient cohort and independent prospective validation are needed to firmly establish the clinical utility of cfChIP-seq.

Recent studies have described methylation-based methods for subtyping SCLC preclinical models and patient samples^62,63^. However, unlike cfChIP-seq, which directly reflects the tumor transcriptional profile, the relevance of plasma methylation patterns to SCLC subtypes and their representation of the tumor’s transcriptional landscape remains less clear. The direct link between chromatin state and gene expression strengthens the utility of cfChIP-seq, suggesting its potential to uncover additional transcriptional characteristics of tumors. In contrast, methylation-based approaches may require the development of specific classifiers for relevant features, limiting their immediate clinical applicability.

This study also provides a broadly applicable framework for benchmarking features of cfDNA against the tumor molecular profile. While plasma cfChIP-seq shows good agreement with tumor RNA-seq, the correspondence is imperfect due to multiple factors. First, there are inherent differences between chromatin state, which is to a large extent on or off, and gene expression, which has a large dynamic range ^29,64,65^. In addition, several factors confound our estimations of both the plasma and tissue compartments. Plasma cell-free chromatin reflects contributions from multiple sources, which also include tumors. Tumor RNA-seq contains contributions from multiple sources in addition to the tumor including tissue-infiltrating immune cells, stromal cells, endothelial cells, and more. Thus, to understand the plasma-tumor correspondence, we need to account for the different cell types contributing to each compartment. By explicitly accounting for differences in SCLC-score we could extract tumor-specific and tumor-extrinsic features even when the tumor contribution is low. Ideally, we would deconvolve the signal from these different sources and extract from plasma samples information about the state of multiple cell types (e.g., tumors, immune cells, stroma cells). Establishing reliable deconvolution requires better references of the ChIP-seq profiles of tumor cells and biopsies, together with baseline estimates of their fractions in each sample^31^.

Overall, our study bridges the gap between molecular studies and patient treatment in SCLC, an exceptionally lethal malignancy, which to date is treated as a homogenous disease with identical treatments for all patients. Moreover, this work suggests the applicability of cfChIP-seq to a wider context to profile and subtype tumors, in a way that can be transformative for patient care across multiple cancer types.

## Materials and Methods

### Patients

We undertook an observational study using plasma collected from patients with small cell cancer who received care at the National Cancer Institute (NCI). Patients were enrolled in therapeutic clinical trials (ClinicalTrials.gov #NCT02484404; NCI protocol #15-C-0145; ClinicalTrials.gov #NCT02487095; NCI protocol #15-C-0150; ClinicalTrials.gov #NCT02769962; NCI protocol #16-C-0107; ClinicalTrials.gov # NCT03554473; NCI protocol #18-C-0110; and ClinicalTrials.gov # NCT03896503; NCI protocol #20-C-0009). We also collected samples from small cell cancer patients who were enrolled in the NCI thoracic malignancies natural history protocol (ClinicalTrials.gov # NCT02146170; NCI protocol #14-C-0105). See **Supplementary table 1** for information per patient. If tumor samples were available, we also sequenced their RNA at matched or different time points of when the plasma was collected. The human subjects committee at NCI approved the studies; all patients provided written informed consent for plasma, tumor, and matched normal sample sequencing.

### Tumor RNA sequencing and Immunohistochemistry

Formalin-Fixed, Paraffin-Embedded (FFPE) tumor tissue samples or frozen tumor samples in selected samples were prepared for RNA-seq. RNA enrichment was performed using TruSeq RNA Exome Library Prep according to the manufacturer’s instructions (Illumina, San Diego). Paired-end sequencing (2 x 75 bp) was performed on an Illumina NextSeq500 instrument. RNA was extracted from FFPE tumor cores using RNeasy FFPE kits according to the manufacturer’s protocol (QIAGEN, Germantown, MD). RNA-seq libraries were generated using TruSeq RNA Access Library Prep Kits (TruSeq RNA Exome kits; Illumina) and sequenced on NextSeq500 sequencers using 75bp paired-end sequencing method (Illumina, San Diego, CA). Reads were aligned to the human genome (hg19) using STAR (2.7.10a)^66^. For transcriptomic analyses, raw RNA-Seq count data were normalized for inter-gene/sample comparison using TMM-CPM, as implemented in the edgeR R/Bioconductor package^67^. Immunohistochemistry for INSM1 was done as we previously described^21^.

### Tumor volumetric estimation

#### cfDNA Tfx

The DSP Circulating DNA kit from Qiagen was utilized to extract cell-free DNA from aliquots of plasma which were eluted into 40-80 uL of re-suspension buffer using the Qiagen Circulating DNA kit on the QIAsymphony liquid handling system. Library preparation utilized the Kapa Hyper Prep kit with custom adapters (IDT and Broad Institute). Samples were sequenced to meet a goal of 0.1x mean coverage utilizing the Laboratory Picard bioinformatics pipeline, and Illumina instruments were used for all cfDNA sequencing with 150 bp and paired-end sequencing. Library construction was performed as previously described ^68^. Kapa HyperPrep reagents in 96-reaction kit format were used for end repair/A-tailing, adapter ligation, and library enrichment polymerase chain reaction (PCR). After library construction, hybridization and capture were performed using the relevant components of Illumina’s Nextera Exome Kit and following the manufacturer’s suggested protocol. Cluster amplification of DNA libraries was performed according to the manufacturer’s protocol (Illumina) using exclusion amplification chemistry and flowcells. Flowcells were sequenced utilizing Sequencing-by-Synthesis chemistry. Each pool of whole genome libraries was sequenced on paired 76 cycle runs with two 8 cycle index reads across the number of lanes needed to meet coverage for all libraries in the pool. Somatic copy number calls were identified using CNVkit^69^ (version 0.9.9) with default parameters. Tumor purity and ploidy were estimated by sclust^69,70^ and sequenza^71^. cfDNA Tfx was estimated based on the somatic copy number alteration profiles using ichorCNA, a previously validated analytical approach^33,71^.

#### CTCs

CTCs were detected from 10 mL of peripheral blood drawn into EDTA tubes. Epithelial cell adhesion molecule (EpCAM)-positive CTCs were isolated using magnetic pre-enrichment and quantified using multiparameter flow cytometry. CTCs were identified as viable, nucleated, EpCAM+ cells that did not express the common leukocyte antigen CD45, as described previously^20^.

#### Radiological volumetric segmentation

We performed volumetric segmentation, a three-dimensional assessment of computed tomography as previously described ^72^. that may more accurately predict clinical outcomes than conventional evaluation by Response Evaluation Criteria in Solid Tumors (RECIST). Briefly, experienced radiologists reviewed the computed tomography sequences to determine the best ones to use for segmentation using the lesion management application within PACS (Vue PACS v 12.0, Carestream Health, Rochester, NY). We also assessed sizes of lesions by RECIST v.1.1.

### Plasma cfChIP-seq

#### Sample collection

Plasma for cfDNA was collected in EDTA tubes and centrifuged at 4C at 1500xg for 10 minutes. Plasma was carefully transferred into Eppendorf tubes, centrifuged a second time at 4C at 10,000xg for 10 minutes, and transferred into standard cryovials.

#### Bead preparation

50μg of antibody was conjugated to 5mg of epoxy M270 Dynabeads (Invitrogen) according to manufacturer instructions. The antibody-beads complexes were kept at 4°C in PBS, 0.02% azide solution.

#### Immunoprecipitation, NGS library preparation, and sequencing

Immunoprecipitation, library preparation, and sequencing were performed by Senseera LTD. using previously reported protocols^27^, with certain modifications that increase capture and signal-to-background ratio. Briefly, ChIP antibodies were covalently immobilized to paramagnetic beads and incubated with plasma. Barcoded sequencing adaptors were ligated to chromatin fragments and DNA was isolated and next-generation sequenced.

#### Sequencing Analysis

Reads were aligned to the human genome (hg19) using bowtie2 (2.3.4.3) with ‘no-mixed’ and ‘no-discordant’ flags. We discarded fragment reads with low alignment scores (-q 2) and duplicate fragments. See **Supplementary Table 3** for read number, alignment statistics, and numbers of unique fragments for each sample.

Preprocessing of sequencing data was performed as previously described^27^. Briefly, the human genome was segmented into windows representing TSS, flanking to TSS, and background (rest of the windows). The fragments covering each of these regions were quantified and used for further analysis. Non-specific fragments were estimated per sample and extracted resulting in the specific signal in every window. Counts were normalized and scaled to 1 million reads in healthy reference accounting for sequencing depth differences.

### ATAC-seq library preparation data processing

Tissue samples for ATAC-seq were directly obtained from patients and PDX’s (patient-derived xenograft) models of different generations. The collected tissue samples were flash frozen and kept at -80 degree Celsius and shipped to Novogene for sequencing. Samples were sequenced at Novogene using NovaSeq 6000 as a sequencing platform with 150bp paired-end sequencing mode. The obtained reads were trimmed using trimmomatic (version 0.39), filtered and aligned to the human genome (hg19) using BWA (version 0.7.17). PCR duplicates and reads mapped to the mitochondrial chromosome or repeated regions were removed.

### Statistical analysis

#### Genes elevated in SCLC compared to healthy plasma

A reference of healthy plasma gene baseline and statistical test to evaluate whether a gene is significantly elevated compared to the baseline were computed as previously described^27^.

#### SCLC score

To estimate SCLC-score reflecting tumor-related fraction in samples, we performed a leave-one-out non-negative least square using the ‘nnls’ R package (1.4).

Denoting by *G* the number of gene promoter, and by *S* the number of samples in the reference cohort, a matrix *X*_*GX*(*S* + 1)_ is composed containing the counts per promoters in the *S* healthy samples, with the addition of a ‘SCLC prototype’ composed on the mean promoter counts of the 10% SCLC samples with most genes significantly higher than healthy (n=15). For every vector *Y*_1*XG*_ of counts per promoter in the SCLC samples, we estimate the non-negative coefficient *β* by computing *argmax_β_* ∥*Xβ* - *Y*∥^2^ subject to *β* ≥ 0. A similar process is computed for healthy samples with the exception that for the *i_th_* sample we use the matrix *X_-i_* where the column containing that sample is eliminated. The estimated *β* is normalized to 1 by computing 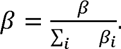 The tumor-related fraction is defined as the value of *β_SCLC_* – the fraction assigned to the SCLC prototype. The healthy fraction of the *i_th_* sample is defined as ∑_*i≠SCLC*_ *β_i_*- the sum of fractions assigned to all other healthy components.

An alternative approach was tested using a reference atlas of roadmap tissue data. In this approach the healthy fraction is set to be the sum of fractions assigned to common tissues observed typically in healthy samples (neutrophils, monocytes, megakaryocytes) and the tumor-related fraction is defined as 1 - the tumor-related fraction. The two approaches resulted in very similar estimations for the vast majority of samples (not shown).

We compute the principal component analysis (PCA) of all gene counts in all SCLC and healthy samples as implemented in the R ‘prcomp’ function. Scree plot was generated using the ‘fviz_eig’ function of the R ‘factoextra’ package (1.0.7).

#### Tissue signatures

Genomic regions selected for tissue-specific markers are as described previously^27^. cfDNA of SCLC patients consists of tumor-related cfDNA above the hematopoietic-derived cfDNA observed in healthy individuals. Estimation of the absolute contribution of tissues to the cfDNA pool was calculated by multiplying the normalized signal per signature and the estimated number of reads in the sequencing library size ^27^ which approximates the cfDNA concentration.

#### Lung single-cell signatures

To define pulmonary cell-type specific signatures, we made use of published pulmonary scRNA-seq data^41^ using cluster-specific marker gene sets defined by the authors. To increase the specificity of cell-type signature in the cfDNA context, we include only genes that meet the following criteria:

a. The percent of cells outside the cluster that express the genes is less than 0.1.
b. The average log fold-change expression in the cluster compared to other clusters is more than 2.
c. The adjusted p-value of the gene is less than 0.1.
d. The mean gene promoter read counts in cfDNA in healthy reference is less than 3.

The last criterion aims to remove from the signature genes that are not lung-specific but appear in circulation as part of normal cell turnover. A summary of the genes included in every lung cell-type signature can be found in **Supplementary Table 5**.

#### ChIP-RNA correlation

To examine the relationship of tumor RNA-seq of tumor and plasma ChIP-seq, we computed the Pearson correlation of all genes across the samples with matching tumor and plasma samples. The correlation was computed between TSS counts of the ChIP-seq and CPM and compared to the correlation achieved from a random permutation of the ChIP-seq samples. The correlation was computed for all samples with matching tumor-plasma (n=73) and separately for samples with SCLC-score > 0.5 (n=21) and matching tumor.

The dynamic range of ChIP-seq gene g was calculated by the 95th percentile - 5th percentile of the normalized reads across all plasma samples. High dynamic range genes were set to be genes with a dynamic range > 20. The dynamic range of the RNA-seq of gene g was calculated similarly for CPM.

##### Subtyping of plasma samples

Based on the RNA-seq distribution, we defined the following CPM cutoffs: *ASCL1* = 10, *NEUROD1* = 60, *POU2F3* = 40 and *ATOH1* = 4. Plasma samples with matching tumors were annotated as TF-positive for each of the four TFs.

##### *ASCL1* specific genomic regions

Tiling of the genome to genomic regions (windows) was performed as previously described^27^. The following procedure was performed to select informative genomic regions. Given a set of labeled samples:

1. Select samples with SCLC score > 0.05
2. Select genomic regions whose mean healthy reference < 0.3 and mean over training sample > 3.
3. Create a matrix of values that represent the 100% tumor contribution to the sample: 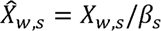 where *X_w,s_* is the observed count at region (window) *w* in sample *s* and *β_s_* is the estimated SCLC score of sample s
4. For each region *w* perform a t-test for the hypothesis that the mean in the positive samples is higher than the negative one.
5. Choose the *k* windows with smallest (most significant) *p*-values

To evaluate different choices of *k*, we performed leave-person-out cross validation, where we removed the samples of one patient from the training set, ran the procedure and evaluated the resulting signature on the held-out samples.

## Acknowledgments

We would like to thank Alon Chappleboim and members of the Friedman lab for their comments and suggestions. We would kindly like to thank Ahmad Shafiei and Mohammad Bagheri from Department of Radiology and Imaging Sciences, Center for Cancer Research, National Cancer Institute, Bethesda, USA for the volumetric measurements.

This work was supported in part by the European Research Council (ERC Adg #101019560 “cfChIP” awarded to N.F.).

## Data availability

Plasma cfChIP-seq normalized promoter coverage and tumor RNA-seq CPM values are available in the Zenodo repository <u>10.5281/zenodo.13766908</u>. UCSC Track hub with cfChIP-seq tracks is available at https://files.cs.huji.ac.il/nirf/track_hubs/SCLC-cfChIP/hub.txt. See also https://genome.ucsc.edu/s/nirfriedman/SCLC%20cfChIP%2Dseq.

## Code availability

All script files used for analyzing data of this manuscript are available on

https://github.com/Nir-Friedman-Lab/SCLC_cfChIPseq.

## Supplementary information

- Supplementary Table 1 - Metadata of SCLC patients, blood samples and biopsies used in this study
- Supplementary Table 2 - Sample category (SCLC, NEC, NSCLC, CRC, healthy) for samples used in this study
- Supplementary Table 3 - Sequencing statistics for samples sequenced in this study
- Supplementary Table 4 - SCLC differential genes
- Supplementary Table 5 - Lung cell-type marker genes used in figure 2F-G and S2C
- Supplementary Table 6 - Liver-specific genes used in figure S3E
- Supplementary Table 7 - genomics regions of the ASCL1 signature used in figure 5

## Supplementary figures

**Fig. S1:**
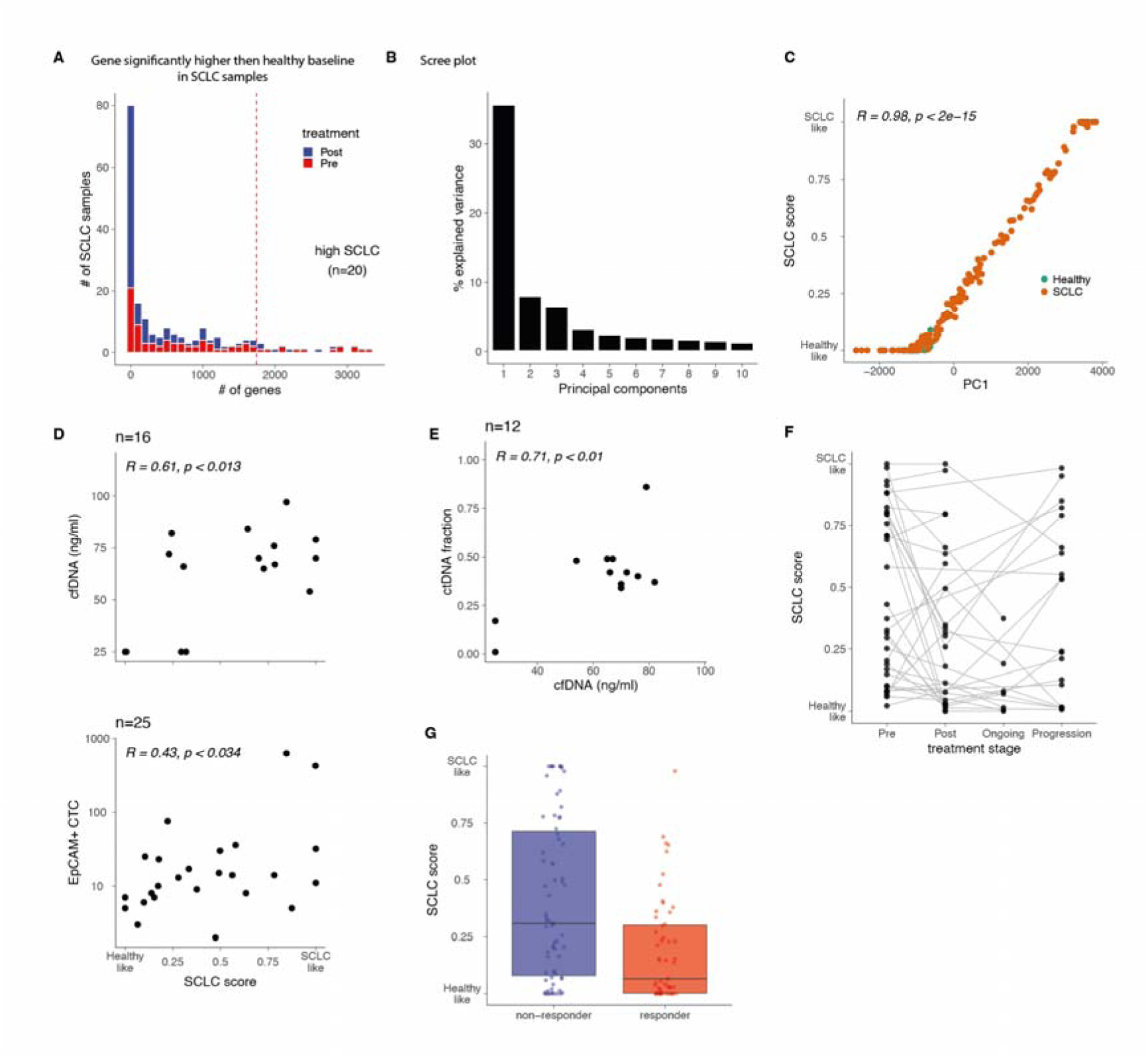
SCLC-score correlates with tumor load and predicts response to treatment. A. Distribution of the number of genes with significantly higher coverage in SCLC samples compared to healthy baseline (Methods). Dashed line represents the 90th percentile. B. Scree plot of PCA (fig. 1D). C. Correlation of PC1 and SCLC-score (presented in fig 1C-D). High correlation indicates that the main variation in the data is driven by the SCLC contribution to the cfDNA. D. Correlation of SCLC-score and other plasma-based methods for tumor load estimations (cfDNA concentration, circulating tumor cells). E. Correlation of cfDNA concentration and ctDNA fraction. F. Dynamics of SCLC-score during treatments of topotecan and berzosertib (an ATR inhibitor) (ClinicalTrial.gov identifier NCT02487095). Gray lines indicate multiple time points of the same individual. G. Median SCLC-score in patients that did or did not respond to various investigational treatments: combination of topotecan and berzosertib (NCT02487095, NCT03896503); olaparib and durvalumab (NCT02484404); nanoparticle camptothecin (CRLX101) and olaparib (NCT 02769962); M7824 (PD-1 inhibitor and TGF-B trapping) and topotecan or temozolomide (NCT03554473). Each dot is a sample, and boxplots summarize distribution of values in each group. Abbreviations: ATR: ataxia telangiectasia and Rad3-related.

**Fig. S2:**
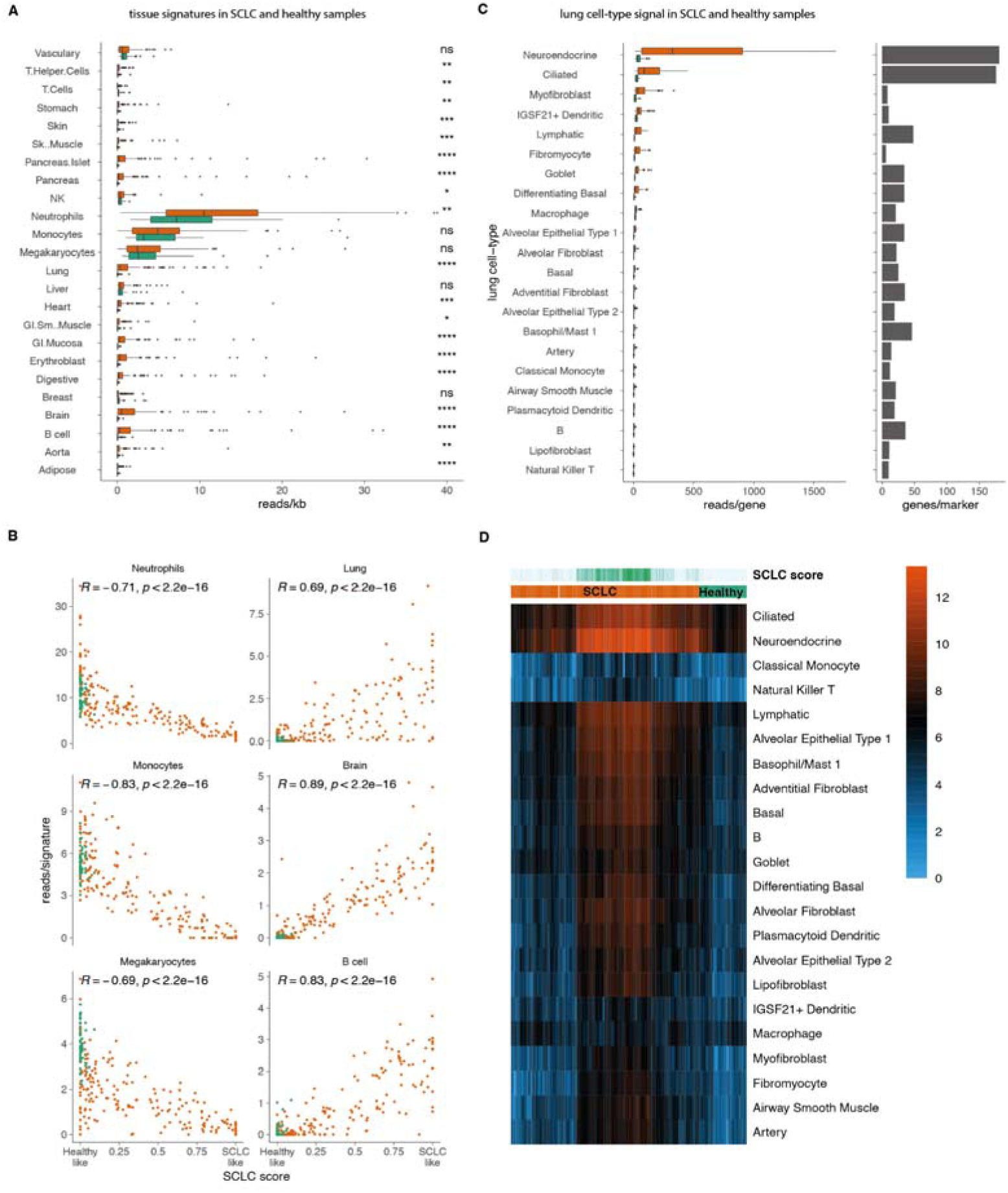
SCLC tissue and cell of origin. A. Distribution of signal for cell-type signatures (same as fig. 2E). B. Correlation of cell-type signature (shown in fig 2E) and SCLC-score. Cell-types observed only in SCLC samples are positively correlated with SCLC-score, while cell-types observed in healthy samples are negatively correlated to SCLC-score. C. Left: number of marker genes used for every cell-type (Methods; **Supplementary Table 5**). Right: Distribution of signal for lung cell-type marker genes (same as fig. 2F). **** and ***: P < 0.0001 and < 0.001, respectively D. Heatmap showing patterns of signal in lung cell-type markers in the SCLC and healthy samples. Color represents log2(1+cumulative coverage in promoters of marker genes).

**figure S3.**
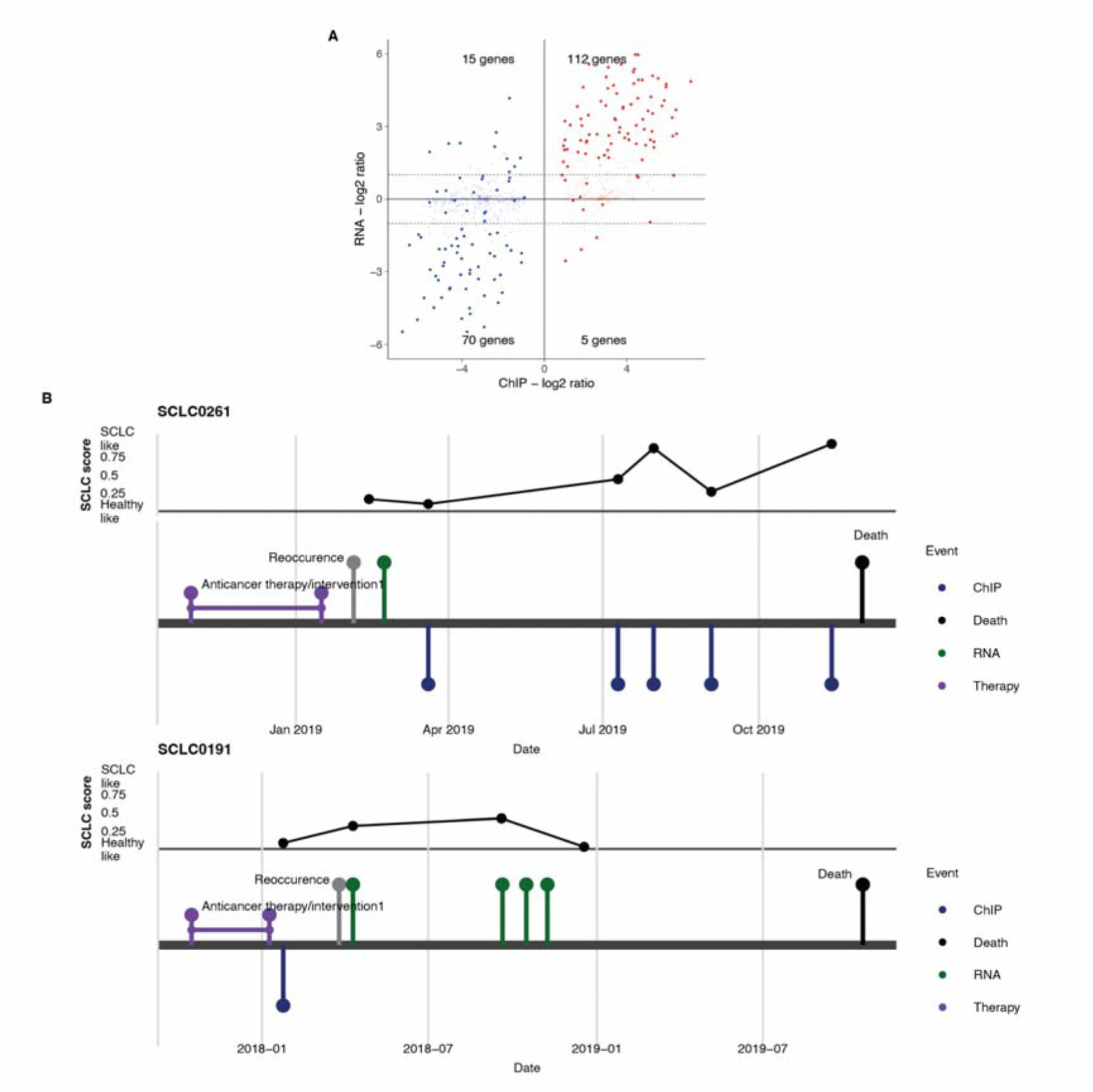
cfChIP and tumor RNA-seq comparison and examples of patient trajectories. A. Comparison of signal ratio of the two individuals shown in Fig. 3A in tumor RNA and plasma cfChIP. Blue and Red points correspond to genes with higher cfChIP-seq counts in every sample. Lighter points represent genes with low RNA counts (combined TMM-CPM< 3). B. Disease time course and interventions (treatment and procedures) of two SCLC patients. Bottom panel - section bordered by purple points represent treatment period, gray and black points represents events of reoccurrence and death, green points represent tumor biopsy RNA-seq samples used in presented in this study. Points below the line represent plasma cfChIP-seq samples presented in this study (green points - samples with time-matched tumor biopsy, red points - samples with unmatched tumor biopsy. Top panel - SCLC score of the cfChIP plasma samples. X-axis represents the dates as in the bottom panel.

**Fig S4:**
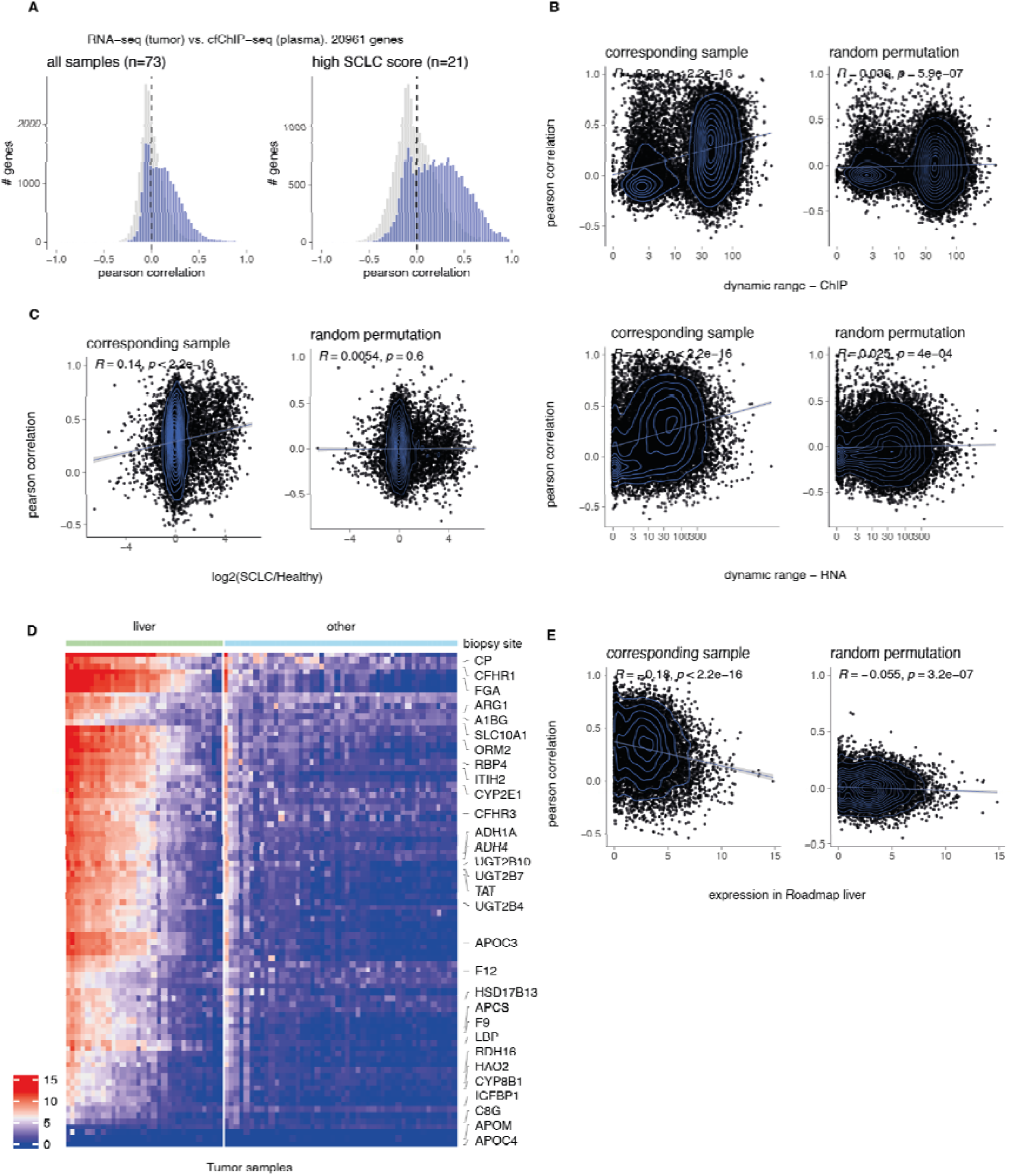
cfChIP-seq RNA correlation confounders. A. Gene level analysis of the correlation in tumor expression level and plasma cfChIP-seq levels (same as in Fig. 3B for all Refseq genes). B. Comparison of the effect of cfChIP-seq and RNA dynamic range (x-axis) on cfChIP-RNA correlation (y-axis) shown in Fig. 3B. Left panel for the observed correlation and right panel for a random permutation (gray histogram of Fig. 3B). Results indicate that genes with low ChIP dynamic range tend to have lower correlation. C. Same as C for SCLC/healthy ratio (x-axis). The ratio was computed using the mean gene counts of high SCLC samples and healthy samples. D. TMM-CPM of tumor samples in liver specific genes. High expression is observed in samples where biopsy was obtained from the liver. E. Same as C for gene expression in the liver (x-axis).

**Fig. S5:**
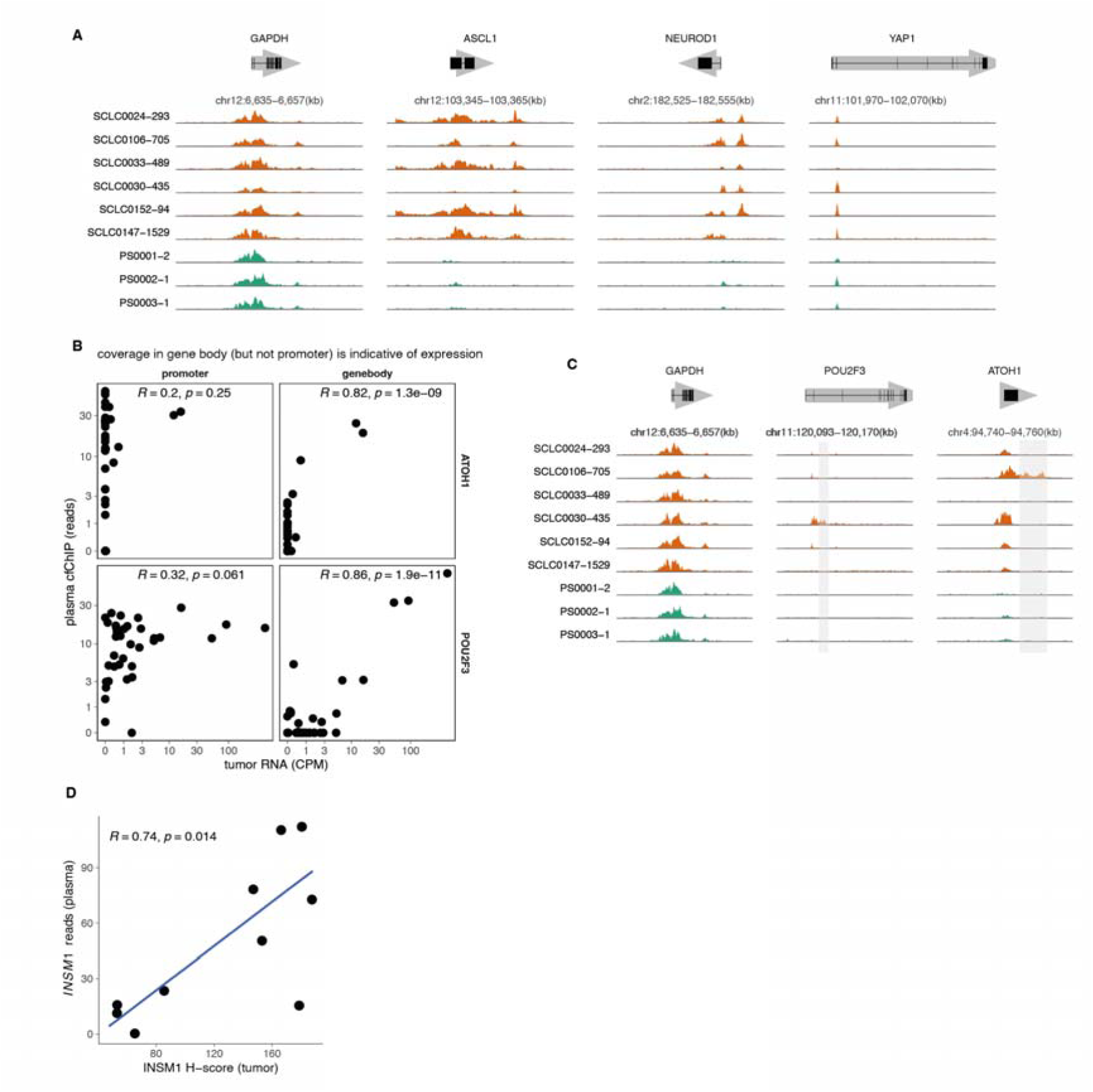
cfChIP signal in SCLC lineage-driving genes. A. Genome browser view of cfChIP-seq signal in SCLC key transcriptional regulators. B. Correlation of plasma cfChIP-seq coverage in promoter and gene body and tumor RNA-seq of the genes *POU2F3* and *ATOH1*. Noticeably, in these genes, the correlation to expression is higher in the gene body. C. Genome browser view of cfChIP-seq signal in panel D. Orange and green tracks represent SCLC and healthy samples respectively. While a signal in the gene promoter is observed in many samples, only in specific samples there is a significant coverage also in the gene body. D. Correlation of plasma cfChIP-seq coverage in the promoter of *INSM1* (y-axis) and the immunohistochemistry H-score of the matching tumors (x-axis).

